# Leveraging the transient statistics of quantal content to infer neuronal synaptic transmission

**DOI:** 10.1101/2024.06.13.598834

**Authors:** Zahra Vahdat, Oliver Gambrell, Jonas Fisch, Eckhard Friauf, Abhyudai Singh

## Abstract

Quantal parameters of synapses are fundamental for the temporal dynamics of neurotransmitter release, forming the basis of interneuronal communication. We propose a class of models that capture the stochastic dynamics of quantal content (QC) - the number of SV fusion events per action potential (AP). Considering the probabilistic and time-varying nature of SV docking, undocking, and AP-triggered fusion, we derive an *exact* statistical distribution for the QC over time. Analyzing this distribution at steady-state and its associated autocorrelation function, we show that QC fluctuation statistics can be leveraged for inferring key presynaptic parameters, such as the probability of SV fusion (release probability), and SV replenishment at empty docking sites (refilling probability). Our model predictions are tested with electrophysiological data obtained from 50-Hz stimulation of auditory MNTB-LSO synapses in brainstem slices of juvenile mice. Our results show that while synaptic depression can be explained by low and constant refilling/release probabilities, this scenario is inconsistent with the statistics of the electrophysiological data that show a low QC Fano factor and almost uncorrelated successive QCs. Our systematic analysis yields a model that couples a high release probability to a time-varying refilling probability to explain both the synaptic depression and its associated statistical fluctuations. In summary, we provide a general approach that exploits stochastic signatures in QCs to infer neurotransmission-regulating processes that are indistinguishable from just analyzing averaged synaptic responses.

## I. Introduction

Action potential (AP)-triggered synaptic transmitter release is a hallmark of interneuronal communication. At a fundamental level, this communication is orchestrated via neurotransmitter-filled synaptic vesicles (SVs) that are docked at sites in the active zone of the axon terminal, and the released neurotransmitter upon action potential arrival impacts postsynaptic neuronal activity. The depletion of SVs in response to a high-frequency AP train is counteracted by their replenishment creating a dynamic equilibrium [1], [2]. Recent work has unmasked diverse types of vesicle pools working sequentially or parallelly with heterogeneity among docked SV/docking sites [3]– [11], and this complexity of presynaptic processes critically shapes both the short-term and the long-term dynamics of neurotransmission in response to a train of APs [12]–[18].

While much work treats neurotransmitter release as a deterministic process [19]–[23], these models fail to capture the variability introduced at each trial by the inherent probabilistic nature of SV recruitment to docking sites and neurotransmitter release by AP-triggered exocytosis of SVs [24]–[27]. Moreover, several experimental and computational publications have argued that these stochastic effects are facilitated information flow across a chemical synapse [28]–[36], and fluctuation statistics of evoked PSCs (postsynaptic currents) provides robust estimates of presynaptic model parameters [37], [38]. In our prior work, we have used the formalism of Stochastic Hybrid Systems to develop mechanistic models of neurotransmission investigating how diverse noise mechanisms shape the statistics of SV counts [39], [40] and their corresponding impact on postsynaptic AP firing times [41].

Here we generalize these models to consider probabilistic docking and undocking of SVs at a fixed number of docking sites. Docked SVs represent the readily releasable pool (RRP) of SVs and each AP triggers probabilistic SV fusion and neurotransmitter release. A key feature of the model is that all these probabilities can *vary arbitrarily over time*, thereby capturing diverse response dynamics, including synaptic facilitation and synaptic depression. These transient parameters reflect a variety of physiological processes during high-frequency stimulation, such as buildup in calcium concentrations in axon terminals or depletion of upstream SV recycling pools that lead to reduced recruitment of SVs to docking sites. For this generalized model, we provide an *exact* analytical solution to the *transient statistical distribution* of the quantal content (QC) defined as the number of SV fusion events per AP. When APs arrive deterministically (i.e., at fixed time points) in the presynaptic axon terminals, the transient QC distribution follows a binomial distribution. In contrast, deviations from the canonical binomial behavior occur when APs arrive stochastically.

The statistical dispersion in QC (as quantified by the QC Fano factor, i.e., the variance divided by the mean) is shown to be a monotonically decreasing function of the mean QC, implying higher statistical fluctuations for stronger synaptic depression. Intriguingly, our analysis shows that *different parameter regimes, that are otherwise indistinguishable from their average QC dynamics, yield contrasting QC Fano factor profiles*. Hence, systematic investigation of transient QC fluctuation statistics from electrophysiological data can be a valuable tool to infer processes regulating neurotransmission. Finally, we investigate the extent of steady-state QC fluctuations as a function of model parameters and also derive an exact analytical expression for the QC auto-correlation function (i.e., the Pearson correlation coefficient between two QCs separated by a given number of stimuli).

The applicability of our modeling results is illustrated by single-cell recordings of QCs for 3,000 stimuli (50-Hz stimulation for 1 min) at the inhibitory glycinergic MNTB-LSO synapses in the auditory brainstem. Classical parameter estimation approaches, such as the method of Elmqvist and Quastel (EQ) [42], or simply fitting the synaptic depression and steady-state performance, predict low constant values for refilling probability and release probability. However, these parameters significantly overestimate the magnitude of statistical fluctuations in QC and anticorrelations between successive QCs as seen in the electrophysiological data. The combination of our analytical results with the experimentally observed fluctuation statistics reveals a dramatically different picture of high release probability and high refilling probability at these robust auditory synapses involved in sound localization. Both are critical for sustained neurotransmission at high frequency and fidelity. We begin with a detailed description of the stochastic model and highlight key underlying assumptions.

### II. Stochastic formulation of synaptic neurotransmission

We consider that APs arrive in the axon terminal with a given frequency *f* at deterministic times {0, 1*/ f* , 2*/ f* , …}. The stochastic dynamics of QC occur as per the following rules with model parameters summarized in Table I for the reader’s convenience:

**TABLE I:** Parameters used in the synaptic transmission model

| Parameter | Description |
| --- | --- |
| $M$ | Number of docking sites for synaptic vesicles (SVs) in the axon terminal. |
| $f$ | Frequency of action potential (AP) arrival at the active zone of the axon terminal. |
| $n_i$ | Random variable representing the number of docked readily releasable SVs just before the arrival of the $i^{th}$ AP, $i \in \{1, 2, \dots\}$. |
| $p_i$ | The probability of a docking site being occupied by an SV just before the arrival of the $i^{th}$ AP. |
| $\bar{p}$ | The steady-state value of $p_i$. |
| $p_{r,i}$ | The probability per docked SV to fuse and release neurotransmitter upon arrival of the $i^{th}$ AP.<br>Represented as $p_r$ when $p_{r,i}$ is constant across APs. |
| $p_{u,i}$ | The probability per site of a docked SV undocking between the $i^{th}$ and $i+1^{th}$ AP.<br>Represented as $p_u$ when $p_{u,i}$ is constant across APs. |
| $p_{d,i}$ | The probability of an empty site getting reoccupied (or refilled) by an SV between the $i^{th}$ and $i+1^{th}$ AP.<br>Represented as $p_d$ when $p_{d,i}$ is constant across APs. |
| $b_i$ | Random variable representing the quantal content (QC) - the number of SVs fusing upon arrival of the $i^{th}$ AP. |
| $\langle b_i \rangle$ | Mean QC across APs. |
| $FF_i$ | The QC Fano factor (variance divided by the mean) quantifying the statistical dispersion in QC $b_i$. |
| $FF$ | The steady-state value of $FF_i := \lim_{i \rightarrow \infty} FF_i$. |
| $\rho$ | The steady-state Pearson correlation coefficient between successive QCs. |

- There are *M* ∈ {1, 2,…} docking sites in the active zone, and each site can be either empty or occupied by an SV.
- We assume that at the start of stimulation when the first AP arrives, each docking site is occupied with probability *p*_1_.
- Upon arrival of the *i*^*th*^ AP where *i* ∈ {1, 2,…}, each docked SV has a probability *p*_*r*,*i*_ of fusing and releasing the neurotransmitter. Upon SV fusion, the corresponding docking site is assumed to instantaneously transition to an empty state. We refer to *p*_*r*,*i*_ as the *release probability* and this is assumed to be an arbitrary function of *i* reflecting transient changes in its value in response to calcium buildup in the axon terminal.
- In between successive APs *i* and *i* + 1, each empty site can be reoccupied with probability *p*_*d*,*i*_. We refer to *p*_*d*,*i*_ as the *refilling probability* and it is also an arbitrary function of *i*.
- Motivated by recent observations of “transient docking” [43], [44], we also consider the scenario where each occupied site can become empty due to SV undocking with probability *p*_*u*,*i*_ between APs *i* and *i* + 1.
- Sites are assumed to be *identical* in terms of their refilling/undocking/release probabilities and operate *independently* of each other.

We refer the reader to Appendix A, where probabilities *p*_*d*,*i*_ and *p*_*u*,*i*_ are directly linked to the kinetic rates of SV docking and undocking at individual sites. These probabilities are also linked to AP timing and thus change with stimulation frequency A sample realization of the model is shown in Fig.1B.

**Fig. 1:**
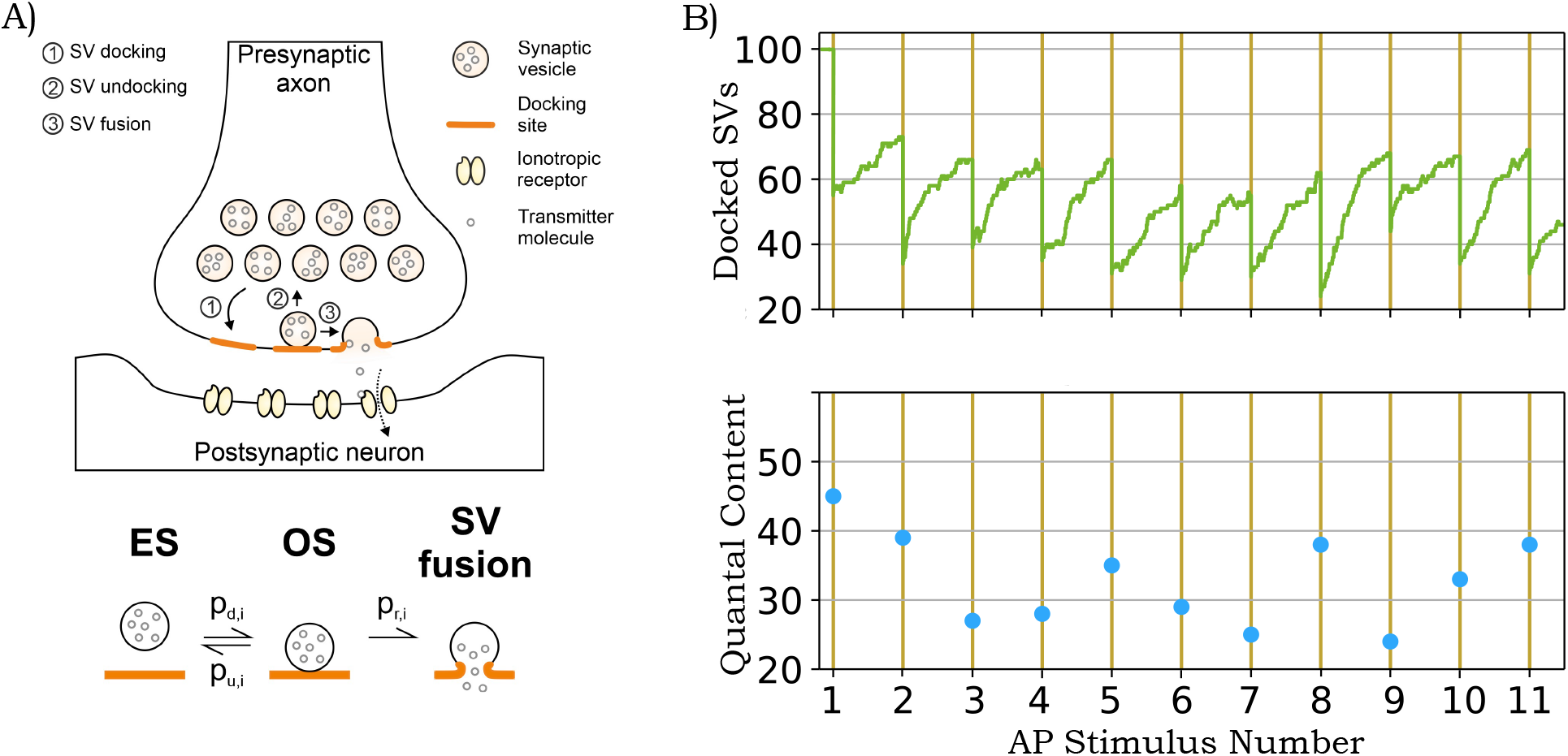
Schematic of a chemical synapse and sample realizations of its corresponding stochastic model. A) The process of SV docking and undocking in the active zone of the axon terminal, and evoked release of neurotransmitter molecules. The lower panel shows the different time-varying probabilities *p*_*d*,*i*_, *p*_*u*,*i*_, *p*_*r*,*i*_ related to the *i*^*th*^ AP where *i* ∈ {1, 2,…}, that govern the reversible transitions between an empty site (ES) and an occupied site (OS) upon SV docking/undocking, and SV fusion (see text for details). B) A sample stochastic realization of the model showing a buildup in the number of docked SVs between successive APs, and a reduction in docked SV numbers from fusion and neurotransmitter release in response to APs (top). The corresponding quantal content (QC) – the number of SV fusion events per AP – is shown in the bottom plot. The parameters here are taken as *M* = 100 with constant release probability *p*_*r*_ = 0.5, refilling probability *p*_*d*_ = 0.4 and undocking probability *p*_*u*_ = 0.1.

## III. Transient distribution of quantal content

Having defined the stochastic model in the previous section, we next first present our main theoretical result quantifying the *exact transient* statistical distribution of QC.

### A. General case of time-varying parameters

Given an initial probability of *p*_1_ of a docking site being filled, a sequence of time-varying probabilities *p*_*r*,*i*_, *p*_*u*,*i*_, *p*_*d*,*i*_ for *i* ∈ {1, 2,…}, then the number of docked SVs *n*_*i*_ just before the arrival of the *i*^*th*^ AP follows the binomial distribution

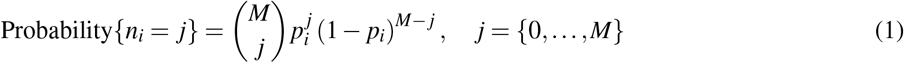

corresponding to each of the *M* sites with probability *p*_*i*_. This probability *p*_*i*_ is the solution to the recursive equation

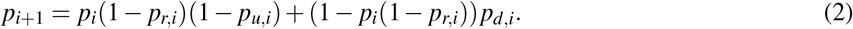

The number of SVs fusing to release neurotransmitter *b*_*i*_ (i.e. the QC) in response to the *i*^*th*^ AP follows the binomial distribution

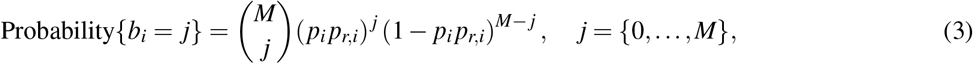

where the binomial coefficient 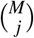 is defined as

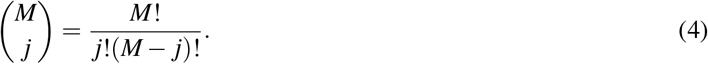

The detailed proof can be found in Appendix B. While this result is for deterministic arrivals of APs, it can be easily generalized to consider the time between APs following an independent and identically distributed (i.i.d.) random variable, in which case *b*_*i*_ no longer follows a binomial distribution (see Appendix C). From this theorem, the average QC across stimuli is given by

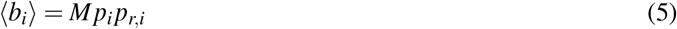

where angular brackets ⟨ ⟩ denote the expected value of random variables and random processes. The statistical dispersion in QC at the *i*^*th*^ stimulus is quantified using the Fano factor

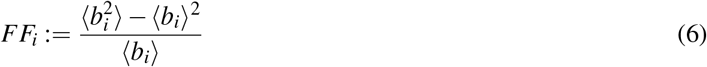

defined as the variance divided by the mean. For a binomial random variable *b*_*i*_

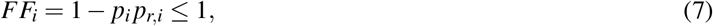

and is always upper-bounded by one. *An important consequence of this result is that one can directly connect the mean QC for the i*^*th*^ *AP to the corresponding statistical fluctuations in QC for any arbitrary time-varying probabilities p*_*r*,*i*_, *p*_*u*,*i*_, *p*_*d*,*i*_. From (5) and (7), one can rewrite the QC Fano factor as

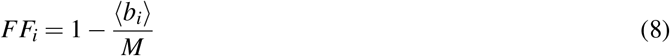

that increases with decreasing ⟨ *b*_*i*_⟩, *implying that a stronger depression in the quantal content is associated with increased randomness in evoked release* (Fig. 2). As ⟨ *b*_*i*_⟩ → 0, *FF*_*i*_ → 1, where a Fano factor of one corresponds to Poisson-distributed *b*_*i*_.

**Fig. 2:**
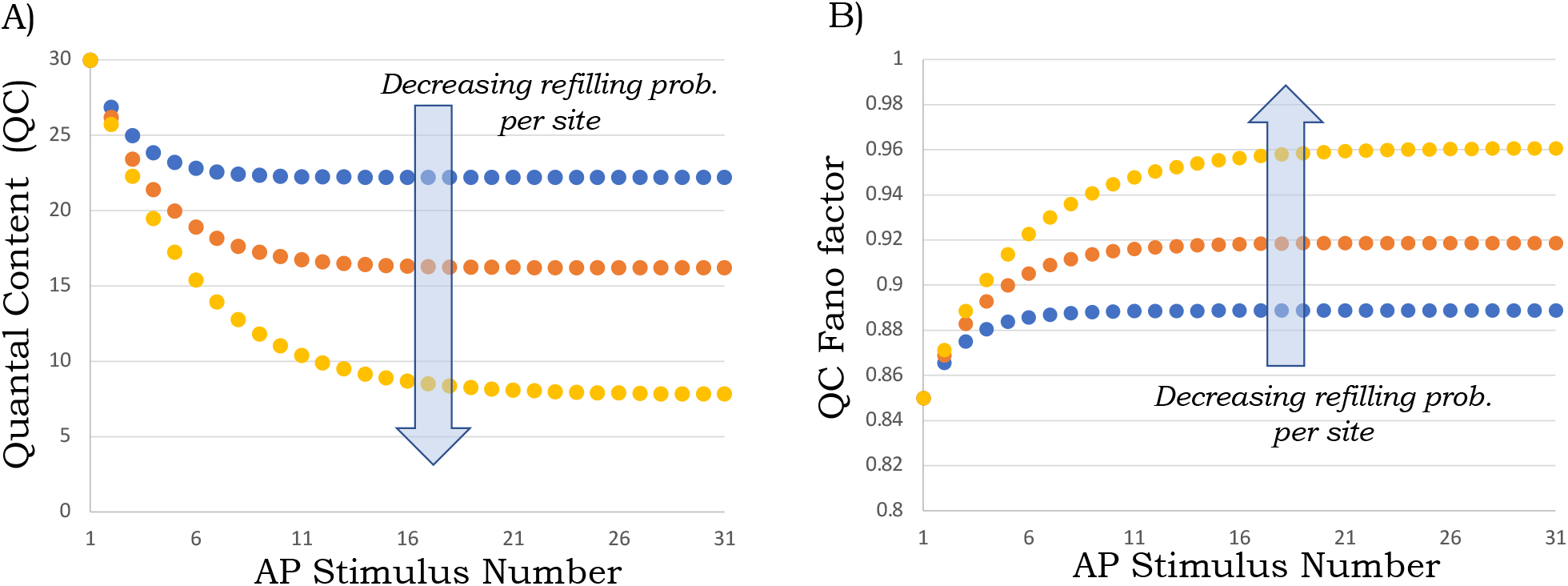
Transient reduction in quantal content (QC) is associated with an increased QC Fano factor. A) The average QC, i.e, the average number of synaptic vesicles (SVs) fusing per stimulus as predicted by equations (5) and (10) for a constant release probability *p*_*r*_ = 0.15, undocking probability *p*_*u*_ = 0. For these plots, the number of docking sites is assumed to be *M* = 200 that are all filled upon arrival of the 1st AP (*p*_1_ = 1). The per-site SV refilling probability is taken to be *p*_*d*_ = 0.3 (blue line), 0.15 (orange line), and 0.05 (yellow line) for the three different lines. B) The corresponding QC Fano factor *FF*_*i*_ over time as predicted by (7).

### B. Special case of time-invariant parameters

An important special case is when refilling/undocking/release probabilities take constant values independent of the AP stimulus number

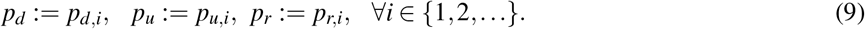

Then, solving the recurrence equation (2) yields

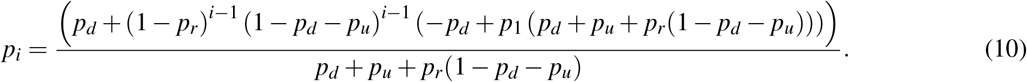

and the corresponding statistical fluctuations in QC are binomially distributed with mean *Mp*_*i*_ *p*_*r*_ and Fano factor 1 − *p*_*i*_ *p*_*r*_. When *p*_*u*_ = 0 and *p*_1_ = 1, (10) reduces to

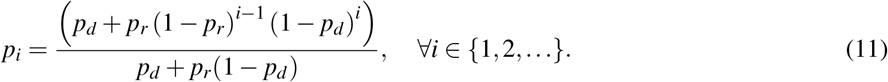

Taking the limit *i* → ∞ in (10) we obtain at steady-state

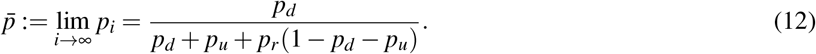

Fig. 2 (left plot) shows the mean QC *Mp*_*i*_ *p*_*r*_ as given by (10), and the depression in the synaptic response is exacerbated with decreasing per site refilling (docking) probability *p*_*d*_. Furthermore, as predicted by (8), the stochasticity in QC increases over time (Fig. 2; right plot).

### C. Indistinguishability of model parameters from mean QC dynamics

It is interesting to point out that the dynamics of the mean QC can be explained by alternative parametric sets that have dramatically different predictions on the QC Fano factor (Fig. 3). To see this, consider the bottom-most yellow trace in the left plot of Fig. 2 that shows the transient reduction in the average QC corresponding to constant and low probabilities *p*_*r*_ = 0.15 and *p*_*d*_ = 0.05. The corresponding Fano factor over time is shown in the left plot of Fig. 2 and also repeated in Fig 3 (top-most curve) for contrasting purposes. Now consider an alternative scenario with a high probability of release. Considering the extreme case of *p*_*r*_ = 1 the average quantal content is given by

**Fig. 3:**
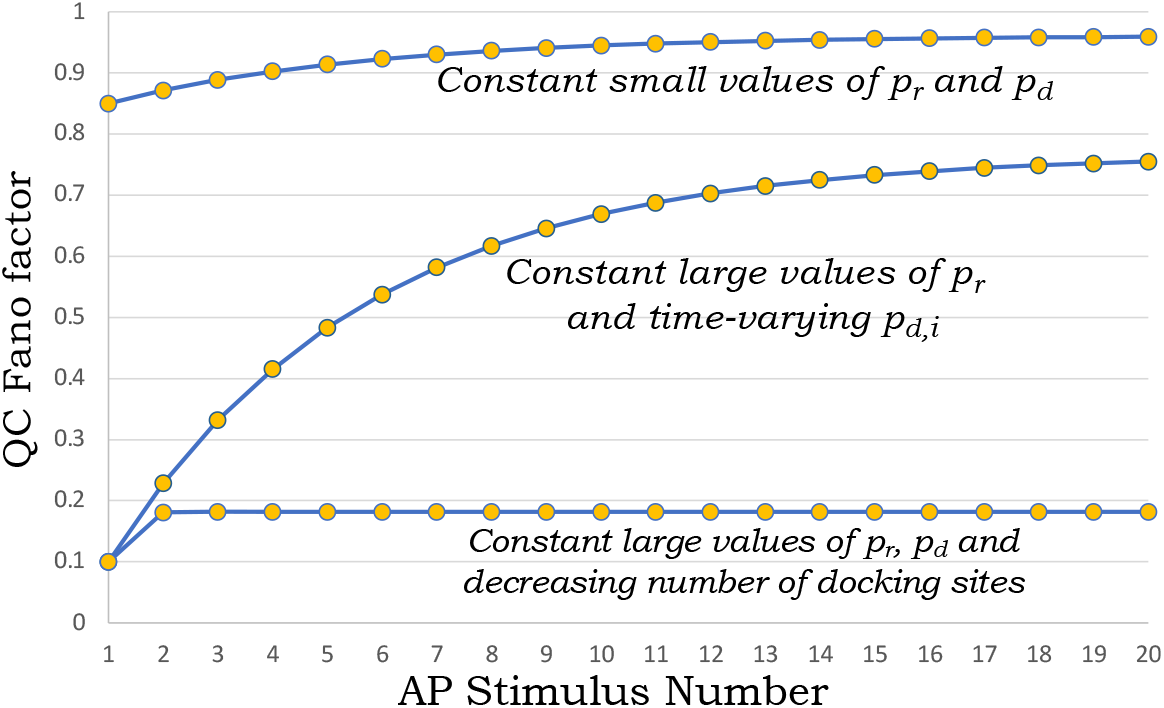
Alternative parameter regimes with identical mean transient QC yield contrasting QC fluctuation statistics. Different predictions for the QC Fano factor, each resulting in the same mean QC corresponding to *p*_*r*_ = 0.15 and *p*_*d*_ = 0.05 in Fig. 2 (the bottom-most, yellow line). The top line corresponds to the Fano factor *FF*_*i*_ predicted for *p*_*r*_ = 0.15 and *p*_*d*_ = 0.05 from (7). The middle curve is obtained from (7) with a high release probability (*p*_*r*_ = 0.9) and a corresponding time-varying refilling probability *p*_*d*,*i*_ to get the same mean synaptic depression. The bottom curve corresponds to (7) with parameters *p*_*r*_ = *p*_*d*_ = 0.9, and here the same mean synaptic depression occurs due to a reduction in the number of docking sites *M*. In all cases, the undocking probability is assumed to be zero (*p*_*u*_ = 0) and each docking site is occupied at the beginning of the challenge period (*p*_1_ = 1).

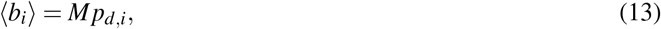

implying that the same decrease in ⟨ *b*_*i*_⟩ can be a result of a decreasing refilling probability *p*_*d*,*i*_ due to depletion of SV pools upstream of the RRP. Note that in this case of *p*_*r*_ = 1 the number of docking sites will have to be much lesser than the former case of *p*_*r*_ = 0.15 and *p*_*d*_ = 0.05 to have the same QC for the first stimulus. However, this scenario predicts a Fano factor profile that starts very low and sharply increases over time (middle-curve in Fig. 3). Finally, consider a third scenario where *p*_*r*_ = *p*_*d*_ = 1 (i.e., all empty sites get occupied in the inter-AP interval and each vehicle is released with probability one upon AP arrival). In this case, the decrease in QC can be potentially explained by a reduction in the number of docking sites *M* due to impaired access to sites. Note that the predicted noise (7) is independent of *M*, and in this case, *FF*_*i*_ is predicted to be low throughout (bottom-most curve in Fig 3). These hypothetical examples serve to emphasize the point that the *QC noise statistics contain useful signatures providing additional insights into the mechanisms underlying synaptic depression*.

## IV. Steady-state qc fluctuation statistics

Assuming that the probabilities *p*_*r*,*i*_, *p*_*u*,*i*_, *p*_*d*,*i*_ reach their respective constant values *p*_*r*_, *p*_*u*_, *p*_*d*_ after several stimuli, we investigate the steady-state QC statistics. To further simplify the formulas we assume that the probability of a docked SV spontaneously undocking during a sustained high-frequency stimulation to be zero (*p*_*u*_ = 0). Our results from the previous section imply that the steady-state QC distribution is binomial with mean *M* 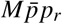 and Fano factor 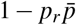 [45], where from (12)

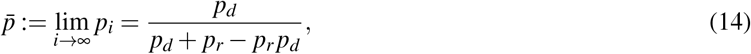

Implying

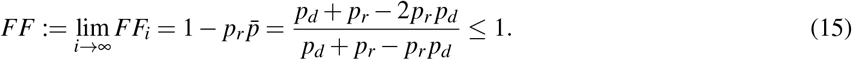

If either of the two probabilities is low (i.e., *p*_*d*_ ≪ 1 or *p*_*r*_ ≪ 1) then *FF* ≈ 1 as illustrated in the left-most-plot of Fig. 4. However high values for both these probabilities result in a low *FF*, with

**Fig. 4:**
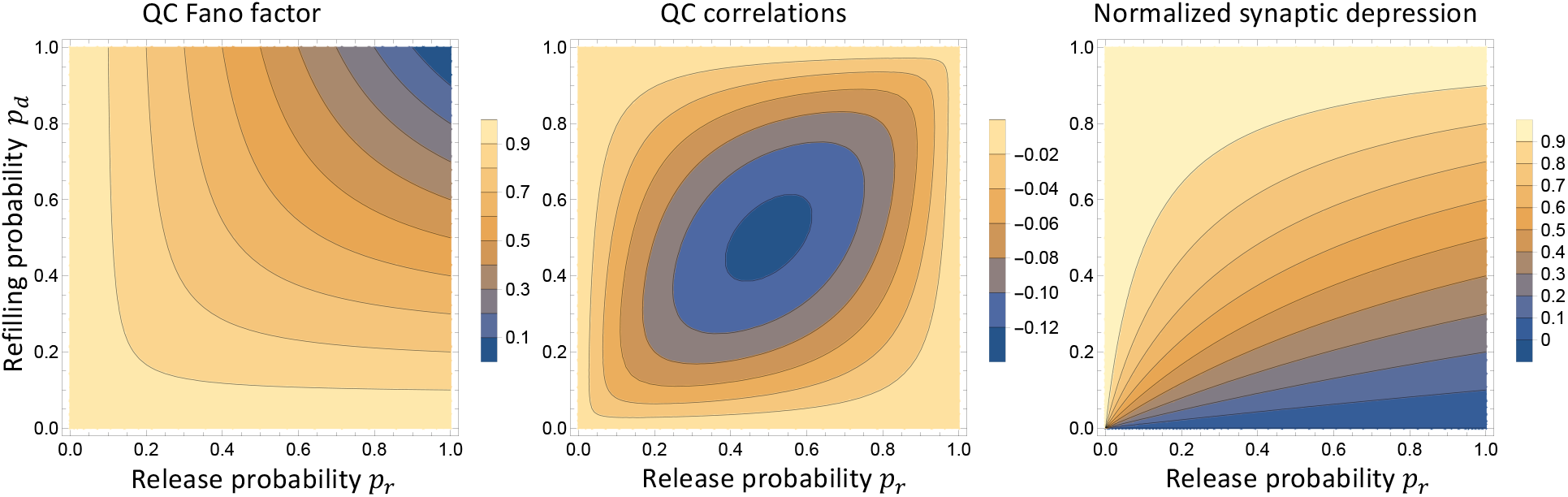
Normalized synaptic depression and QC fluctuation statistics as a function of release and refilling probabilities. *Left*: Plots of the steady-state QC Fano factor as given by equation (15). *Middle*: Steady-state correlation between successive QCs as given by equation (18). *Right*: Normalized synaptic depression assumed to equal to 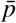 in equation (14) as a function of the release and refilling probabilities. From the middle plot, one can see that if both probabilities *p*_*r*_ and *p*_*d*_ are simultaneously high or simultaneously low, this leads to uncorrelated QCs. However, the two scenarios make contrasting predictions on the Fano factor (left panel). In particular, high probability values lead to a Fano factor close to zero, whereas low values to a Fano factor close to one.

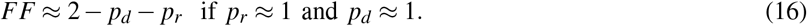

As *FF* in (15) is a function of both probabilities, it by itself cannot be used to infer *p*_*r*_, *p*_*d*_ independently. However, note from (15) that

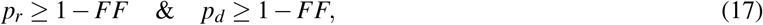

and thus *FF* provides a useful lower bound of both these parameters.

Having determined the steady-state statistical dispersion in QCs, we next focus our attention on correlations between successive QCs. For our stochastic model, the steady-state Pearson correlation coefficient between successive QCs is determined as

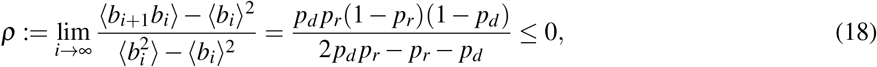

and is predicted to be always non-positive, i.e., a higher-than-average QC would result in the next QC to be lower-than-average due to SV depletion. We refer the reader to Appendix D for proofs and generalization of these results to *p*_*u*_ ≠ 0 and correlations between ⟨ *b*_*i*_⟩ and ⟨ *b*_*i*+*𝓁*_⟩, where *𝓁* ∈ {1, 2,…}. As illustrated in the middle plot of Fig. 4, *ρ* is low if either of the probabilities *p*_*r*_ or *p*_*d*_ takes values close to zero or one, and stronger anticorrelation is seen at intermediate values of both probabilities. The minimum value of *ρ* = −0.125 is attained at *p*_*r*_ = *p*_*d*_ = 0.5. For a given value of *p*_*d*_, *ρ* varies non-monotonically with respect to the release probability (see Fig. S1 in Appendix D) attaining a minimum value when

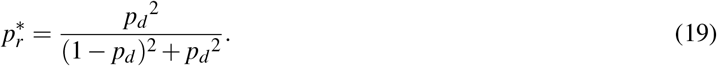

Finally, the right-hand plot of Fig. 4 plots the normalized synaptic depression defined as the steady-state average QC normalized by its corresponding value in response to the first stimulus. This is defined by the ratio

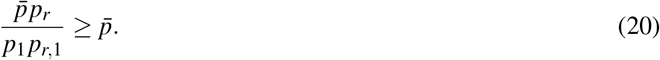

Recall that *p*_1_ is the probability of a docking site being occupied at the first AP, and *p*_*r*,1_ is the corresponding release probability. In many cases *p*_*r*,1_ is much lower than its steady-state value *p*_*r*_ due to calcium buildup in the presynaptic axon terminal, and the ratio (20) is lower bounded by 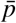 as given by (14). The left-hand plot of Fig. 4 plots the normalized synaptic depression 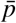 as given by (14) assuming *p*_1_ = 1 and *p*_*r*,1_ = *p*_*r*_, and is sensitive to the refilling probability with

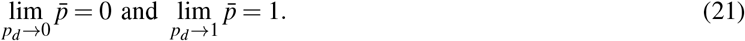

As illustrated next, combining knowledge of normalized synaptic depression and QC fluctuation statistics from electrophysiological data with formulas presented here provides an effective tool to infer model parameters.

## V. MNTB-LSO Synapse: An experimental case study

We apply the mathematical results developed here to the study of neurotransmission in the auditory system. Auditory neurons can fire APs at high rates and must be able to do so continuously to enable sound localization as well as object and speech recognition in noisy environments [46]–[52]. Specifically, we used published data from electrophysiological recordings in juvenile mouse brain slices of the inhibitory glycinergic connection between the medial nucleus of the trapezoid body (MNTB) and the lateral superior olive (LSO) in the medullary brainstem (hereafter referred to as MNTB-LSO synapses). This connection plays a role in sound localization by analyzing interaural intensity differences [53]–[56].

Fig. 5A shows the QC estimations from a whole-cell patch-clamp recording from a single LSO neuron at a 50-Hz challenge for 1 min (3000 stimuli) as taken from [51]. As seen in Fig. 5B, the QC reaches a steady state after an initial decrease (a close-up of Fig. 5A with QC values for the first 20 stimuli). The reader is referred to [51]

**Fig. 5:**
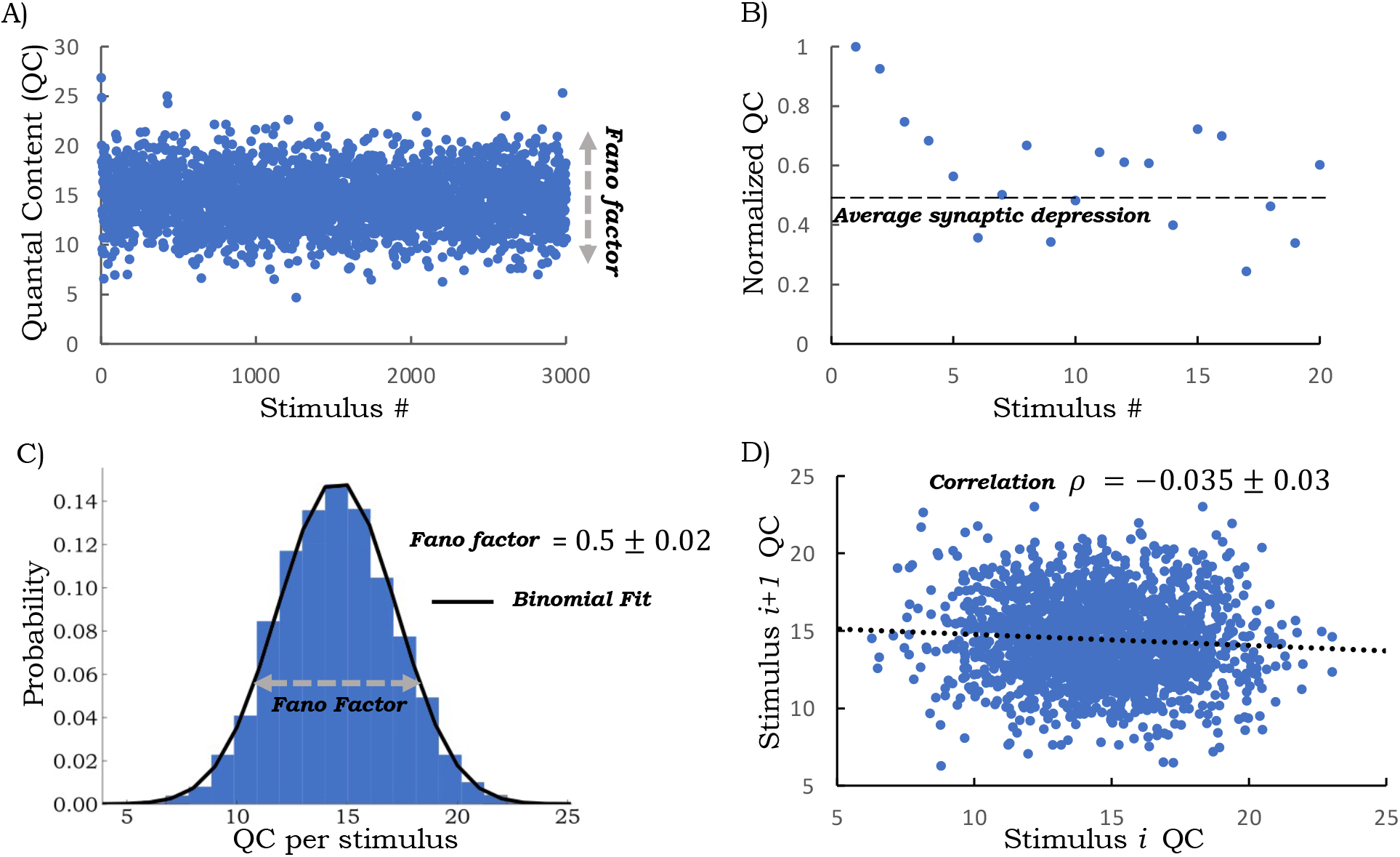
Fluctuation statistics of the quantal content (QC) for the auditory MNTB-LSO synapse. A) Results of a single-cell recording at 50 Hz stimulation for 1 min (3000 stimuli) as obtained from [51]. Each point represents the QC after a single stimulus pulse. B) Normalized QC values to the first 20 stimuli show the initial depression behavior and the subsequent average synaptic depression level. Values are normalized to the first stimulus QC. C) Steady-state QC distribution as obtained using QCs from stimulus numbers 10 to 3000 together with a fit to a binomial distribution. The steady-state QC Fano factor is obtained as *FF* = 0.5 ± .02, where ± denotes the 95% confidence intervals as obtained by bootstrapping. D) The scatter plot between successive QCs from stimulus number 10 to 3000 shows a weak negative correlation with a Pearson correlation coefficient *ρ* = −0.035 ± 0.03.

for experimental details; the QC is obtained by dividing the peak amplitude of evoked PSCs (postsynaptic current) by the average spontaneous PSC peak amplitude in the same neuron. The spontaneous PSCs followed a Gaussian distribution with a mean of 22pA and were found to be the same before stimulation and at the end of the 50-Hz challenge period [51]. Steady-state statistics are quantified using QCs from stimulus number 10 to 3000. We focus on two steady-state metrics: the Fano factor (*FF*; variance divided by mean) and the Pearson correlation coefficient *ρ* between consecutive QCs. This analysis shows *FF* = 0.5 ±.02 with a weak, but statistically significant, negative correlation *ρ* = −0.035 ± 0.03, where ± denotes the 95% confidence intervals as obtained by bootstrapping (Figs. 5C & D).

Assuming constant refilling and release probabilities *p*_*d*_ and *p*_*r*_, respectively, we first fitted the data with the model-predicted average QC over time (normalized by the first stimulus QC) as given by (11)

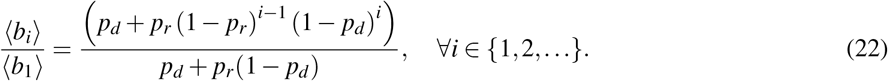

This equation (22) shows a good fit to the synaptic depression observed in the electrophysiological results (Fig. 6A) and results in the inferred values *p*_*r*_ ≈ 0.23 & *p*_*d*_ ≈ 0.2 suggesting that the synapses operate at low values for both these probabilities. Using the classic method of Elmqvist and Quastel (EQ) [42], which assumes no SV replenishment during the first 50 msec of a high-frequency challenge, yields an even lower *p*_*r*_ ≈ 0.12 (using QCs from the first three stimuli). Interestingly, the obtained values for *p*_*r*_ ≈ 0.23 & *p*_*d*_ ≈ 0.2 are incompatible with the steady-state QC fluctuation statistics as reported in Fig. 5. For example, using these values in the mathematical formulas from the previous section demonstrates a much higher model-predicted *FF* of 0.87 and a stronger anticorrelation between QCs (Fig. 6C). To be able to capture these steady-state statistics, one would need *p*_*r*_ ≈ 0.93 & *p*_*d*_ ≈ 0.53 as obtained by simultaneously solving equations (15) and (18) (model-predicted Fano factor and correlation, respectively) with the experimentally determined statistics of *FF* = 0.5 and *ρ* = −0.035. Simulated QCs based on these model parameters are shown in Fig. S3 illustrating the reduction in QC fluctuations for higher values of *p*_*r*_ and *p*_*d*_. These results show that MNTB-LSO synapses operate with a release probability and an SV refilling probability that are much higher than estimated by simply fitting the mean QC dynamics or by using the EQ method.

**Fig. 6:**
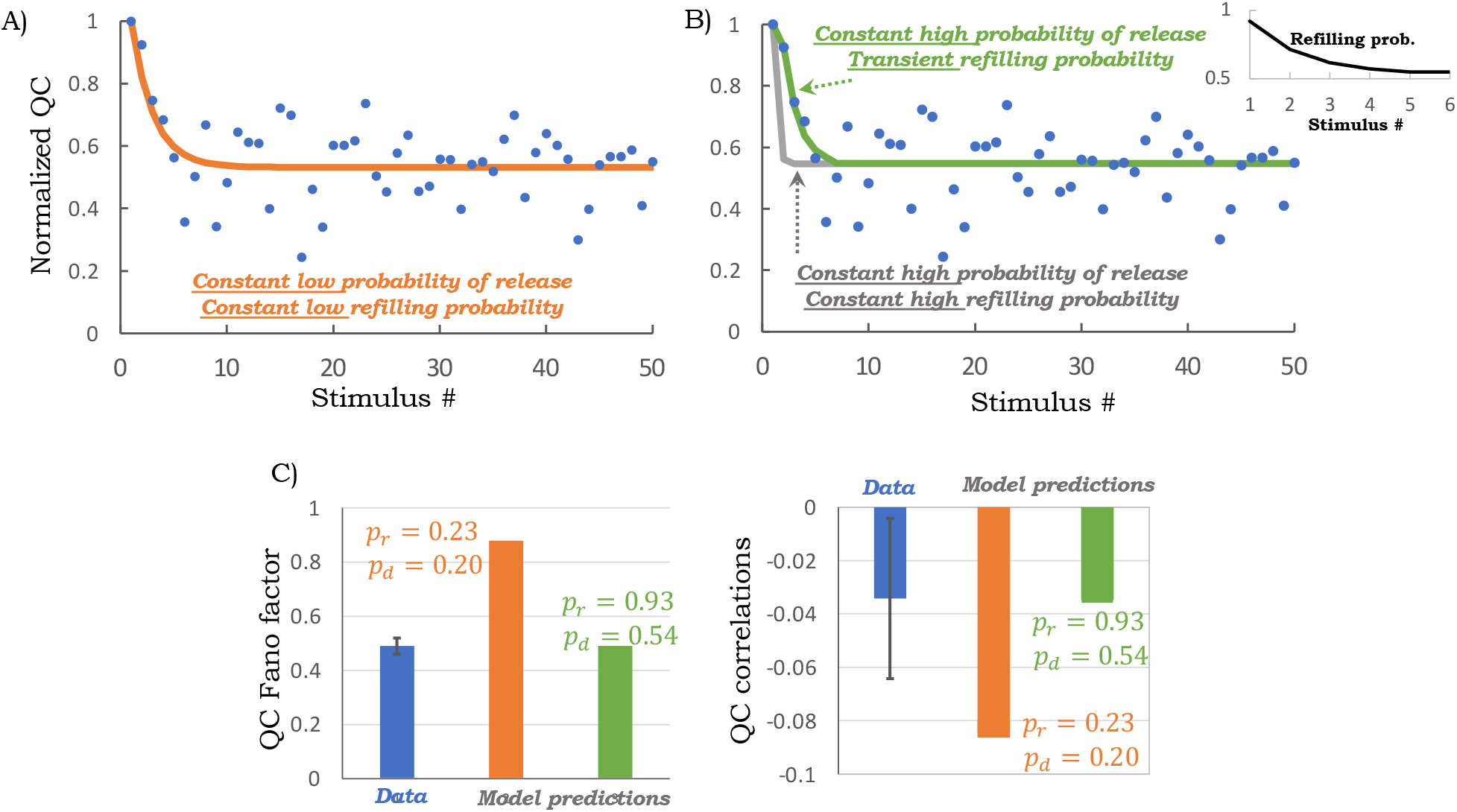
The MNTB-LSO synapses are characterized by high refilling and release probabilities. A) A quantitative fit of equation (22) with *p*_*r*_ = 0.23 & *p*_*d*_ = 0.2 to the transient QC dynamics (orange line). B) Model-predicted QC dynamics as per equation (22) with *p*_*r*_ = 0.93 & *p*_*d*_ = 0.53 (gray line), and as predicted by solving (2) for a constant release probability *p*_*r*_ = 0.93, zero undocking probability *p*_*u*,*i*_ = 0 and a time-varying refilling probability as shown in the inset (green line). C) Model-predicted steady-state QC Fano factors from (15) for constant low probabilities (*p*_*r*_ = 0.23 & *p*_*d*_ = 0.2) or high (*p*_*r*_ = 0.93 & *p*_*d*_ = 0.53) probabilities. Only the latter scenario is consistent with fluctuation statistics from the electrophysiological data. D) Model predicted steady-state correlations between successive QCs from (18) for constant low and high probabilities. Error bars on the data are the 95% confidence interval on the steady-state statistics as obtained from bootstrapping QCs from stimulus numbers 10 to 3000.

Fig. 6B shows the predicted transient dynamics for these high constant probabilities *p*_*r*_ ≈ 0.93 & *p*_*d*_ ≈ 0.53. With these high values, the initial transient is much faster than the data (gray line in Fig. 6B). Our model fits further show that this initial transient is captured by a time-varying refilling probability that starts high at a *p*_*d*_ ≈ 0.95 and reaches its steady-state value of 0.53 within the first five stimuli (see inset of Fig. 6B). In summary, a transient refilling probability coupled to a high release probability explains both the synaptic depression characteristics and the steady-state QC fluctuation statistics (Fig. 6B, green line).

## VI. Discussion

In this contribution, we have investigated the stochastic dynamics of neurotransmission as governed by the depletion and replenishment of a *single* RRP of SVs in response to an AP train (Fig. 1). The model is defined by a fixed number of docking sites *M*, where each site is characterized by three time-varying probabilities:

- The probability *p*_*d*,*i*_ of an empty site becoming docked by a release-ready SV in the inter-spike interval. This probability is monotonically related to the time-varying kinetic rate of SV recruitment to empty sites (Appendix A).
- The probability *p*_*u*,*i*_ of an occupied site becoming empty in the inter-spike interval due to SV undocking or a spontaneous release event.
- The probability *p*_*r*,*i*_ of AP-triggered SV fusion and neurotransmitter release.

We emphasize that the *sites operate independently and are identical in their parameters*. Our main result is: For a deterministic AP train with an inter-spike interval 1 / *f* , where *f* is the frequency of stimulation, the number of release-ready docked SVs just before the *i*^*th*^ AP is binomial with parameters *M* and *p*_*i*_ – each of the *M* sites occupied with probability *p*_*i*_, where *p*_*i*_ is given as the solution to (2). Moreover, the transient QC distribution given by equation (3) is also binomial with parameters *M* and *p*_*i*_ *p*_*r*,*i*_. As discussed further in Appendix C, this result can be generalized to scenarios where the inter-spike interval is random, in which case the QC distribution is non-binomial, and the QC Fano factor can exceed one.

Because of the transient distribution, for any stimulus within an AP train, the QC Fano factor for the *i*^*th*^ AP can be directly related to the corresponding average QC (equation (8) and Fig. 2). Furthermore, alternative parameter regimes leading to the same average QC dynamics can be distinguished by their transient Fano factor profiles (Fig. 3). The above assumption of identical sites can be relaxed by considering another set of sites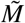 , with different parameters 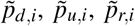. With two classes of docking sites (possibly resulting from differences in their proximity to calcium channels) – *M* sites with parameters {*p*_*d*,*i*_, *p*_*u*,*i*_, *p*_*r*,*i*_}, and 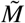 sites with parameters 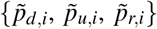– the transient QC is a sum of two binomially-distributed random variables *b*_*i*_ and 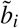 as given by (3) for their respective parameters. Thus, our analytical results can be generalized to consider heterogeneous SV pools operating in parallel, thus resulting in richer neurotransmission dynamics.

By further investigating the process during steady-state transmission, we derive formulas for the Fano factor *FF* and correlation *ρ* between successive QCs as a function of the steady-state refilling and release probabilities *p*_*d*_ and *p*_*r*_, respectively. While high (*p*_*d*_, *p*_*r*_ ≈ 1) as well as low (*p*_*d*_, *p*_*r*_ ≪ 1) values of these probabilities lead to uncorrelated QCs (see the upper-left and lower-right regions of the middle plot in Fig. 4), these regions yield contrastingly different Fano factors, namely a *FF* close to zero when both probabilities are high, yet a *FF* close to one when both probabilities are low. Furthermore, intermediate value of these probabilities (*p*_*d*_, *p*_*r*_ ≈ 0.5) leads to the most anticorrelated QCs (Fig.4). While the theoretical analysis in Section IV considered a zero undocking probability when *p*_*u*_ ≠ 0

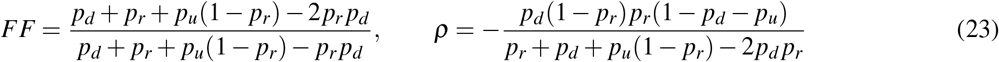

and increasing *p*_*u*_ increases both the *FF* (i.e., higher QC statistical fluctuation) and *ρ* (i.e., more uncorrelated QCs).

Using *FF* and *ρ* from electrophysiological data one can estimate *p*_*d*_ and *p*_*r*_ by simultaneously solving the nonlinear equations (15) and (18). For example, using a *FF* = 0.5 and *ρ* = −0.035 results in two sets of symmetric solutions: 1) *p*_*r*_ = 0.93 & *p*_*d*_ = 0.53 or 2) *p*_*r*_ = 0.53 & *p*_*d*_ = 0.93. The reason is that both (15) and (18) are themselves symmetric with respect to both *p*_*d*_ and *p*_*r*_. In this case, given the high refilling probability, the second solution (*p*_*r*_ = 0.53 & *p*_*d*_ = 0.93) predicts a normalized synaptic depression of 0.96 (steady-state mean QC normalized by the first stimulus). This is inconsistent with the synaptic depression observed in the data, which is closer to 0.55 (Fig. 5). Thus, the first solution (*p*_*r*_ = 0.93 & *p*_*d*_ = 0.53) provides the physiologically relevant parameters that are consistent with the electrophysiological data in all three metrics: normalized synaptic depression, steady-state QC Fano factor, and QC correlation.

The process of narrowing down the feasible parameter space is illustrated in Fig. 7. Given the errors in quantifying QC from evoked PSCs, and other physiological sources of variation, such as differences in AP duration/amplitude [57], the *FF* obtained from electrophysical data is an upper bound on the true *FF* of QC. To this end, one can mark a feasible parameter space consistent with *FF* ≤ 0.55, where the conservative upper bound of 0.55 comes from taking a 10% range around the experimentally-observed *FF* ≈ 0.5 in Fig. 5C. The low *FF* value restricts the parameter space to the upper left corner (i.e., high values of *p*_*d*_ and *p*_*r*_) in the left-most plot in Fig. 7. This can also be seen mathematically in equation (17) where 1 −*FF* provides a lower bound on both *p*_*d*_ and *p*_*r*_: the lower the *FF*, the higher the probabilities. While the results shown in this paper are based on a recording from a single postsynaptic LSO neuron, the analysis of 15 such recordings from in Appendix E shows an *FF* ≤ 0.5, in 80% of the cases (12 of 15), with some MNTB-LSO connections displaying a Fano factor as low as 0.2 (Fig. S2).

**Fig. 7:**
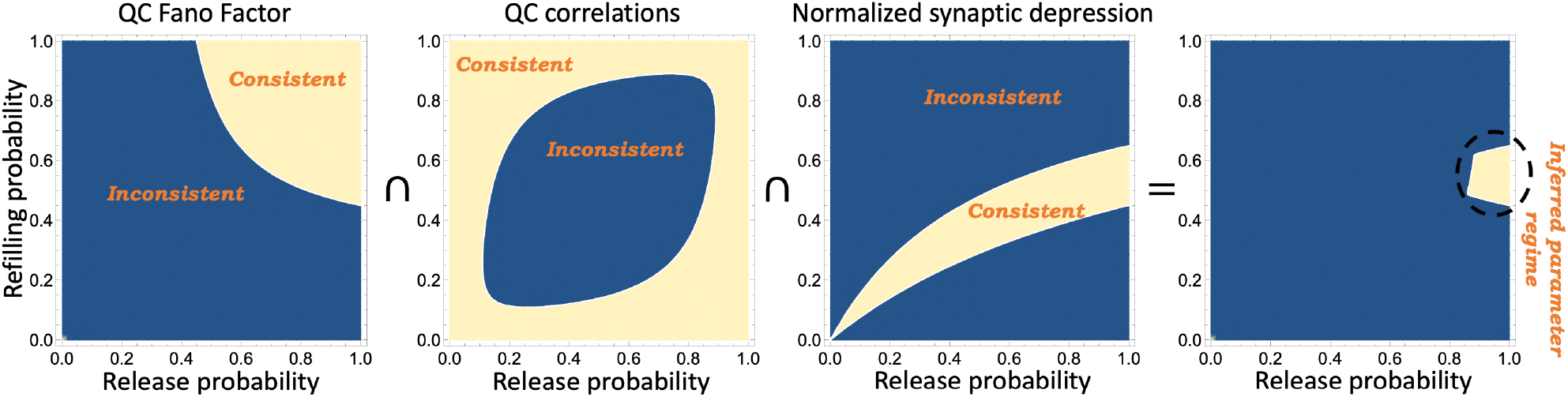
Identification of release and refilling probabilities for MNTB-LSO synapse using QC fluctuation statistics. The left-most-plot marks the region of parameter space (in terms of the refilling and release probabilities *p*_*d*_ and *p*_*r*_, respectively) consistent with a steady-state QC Fano factor as predicted by the formula (15) to be less than *FF* ≤ 0.55. The *FF* upper bound is based on a 10% range around the experimentally-observed *FF* ≈ 0.5 in Fig. 5C. The other plots mark the parameter space consistent with QC correlations as given by (18) to be *ρ* ≥ −0.06 and the normalized synaptic depression as given by (14) in the range 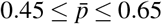 based on the electrophysiological data in Fig. 5. The right-most-plot shows the intersection of all three consistent regions narrowing the parameter space to a region with a high release probability (*p*_*r*_ ≥ 0.85) and 0.45 ≤ *p*_*d*_ ≤ 0.65.

By marking similar regions consistent with the observed QC correlations and normalized synaptic depression in Fig. 7, and taking an intersection of these regions, the probabilities are constrained to the middle-left region of the parameter space (right plot in Fig. 7), corresponding to a high release probability and a refilling probability in the range of 0.53 ± 0.1. To estimate the SV replenishment rate from the refilling probability we use equation (25) with an inter-AP interval of 20 msec and *k*_*u*,*i*_ = 0 resulting in a recruitment rate of 37 SVs per empty site per sec. In this estimation procedure, we have assumed the undocking probability *p*_*u*_ to be zero. For example, for *p*_*u*_ = 0.2, solving equations (23) yields a refilling probability *p*_*d*_ ≈ 0.57 which results in recruiting 54 SVs per empty site per sec. Finally, our modeling indicates that the initial transient depression in QCs is a result of a decreasing refilling probability (inset in Fig. 6B). We plan to capture this phenomenon more mechanistically in future work by considering an SV pool that feeds into the docked, readily releasable pool.

In summary, the exact analytical solution for the transient QC distribution provides an elegant, novel framework to infer presynaptic model parameters from QC fluctuation statistics. We are currently working in close collaboration to test model predictions at the MNTB-LSO synapse for different frequencies and challenge durations. We also plan to investigate other auditory synapses, such as glutamatergic calyces of Held and CN-LSO synapses [51], as well as synapses in the cerebellum [11], [58] and the hippocampus [3]. On a theoretical level, we aim to extend these models to include loosely vs. tightly docked SVs as has been recently reported [59], [60] and consider a repair period for docking sites before they become available for SV refilling [61]. Other avenues of future work involve exploring feedback control of neurotransmission, such as regulating of presynaptic processes by secreted neurotransmitters through “autoreceptors” [62]–[66], and investigate stochastic dynamics of interconnected neurons starting with simple feedforward motifs [67], [68].

## Appendix A

Let *k*_*d*,*i*_ and *k*_*u*,*i*_ be constant kinetic rates with which SVs dock and undock to each docking site, respectively, between the *i*^*th*^ and *i* + 1^*th*^ AP. Then, the probability *x*(*t*) of an empty site to be occupied in the subsequent time *t* evolves as

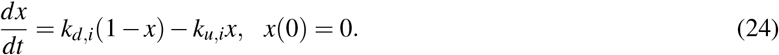

This results in the following refilling probability in time interval *T*_*i*_ between successive Aps

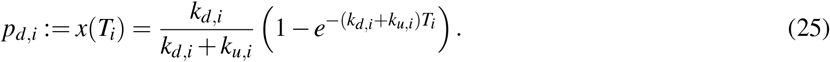

Similarly, solving (24) with initial condition *x*(0) = 1 gives the probability of a docked SV undocking in time interval *T*_*i*_

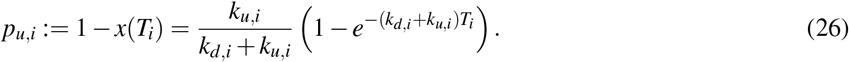

## Appendix B

A detailed proof can be found in the preprint [69] and is also provided here for convenience. We use the notation

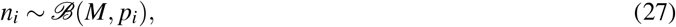

to denote that the random variable *n*_*i*_ follows the binomial distribution with probability mass function (1) that is defined by *M* (the number of trials) and *p* _*i*_ (the success probability in each trial). The probability generating function (pgf) of *n*_*i*_ is given by

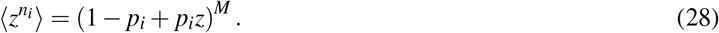

Conditioned on *n*_*i*_, the QC is also binomial *b*_*i*_ ∼ ℬ(*n*_*i*_, *p*_*r*,*i*_), then *b*_*i*_ is itself binomially distributed with parameters

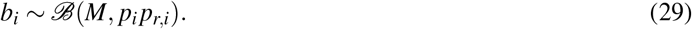

To see this we first find

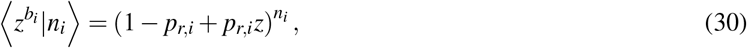

and then unconditioning with respect to *n*_*i*_

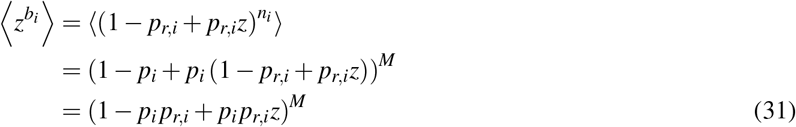

yields the pgf of *b*_*i*_ that is the same as that of a binomially distributed random variable with parameters *M* and *p*_*i*_ *p*_*r*,*i*_.

To prove (27), where *p*_*i*_ is the solution to (2), we use the method of induction, where we know (27) is true for *i* = 1. Now assuming it to be true for any arbitrary stimulus number *i* we show it is also true for *i* + 1. Given *n*_*i*_ ∼ ℬ (*M, p*_*i*_), then the number of SVs 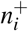 just after the *i*^*th*^ AP will be

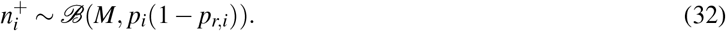

binomially distributed corresponding to each of the docked SVs not releasing with probability 1 − *p*_*r*,*i*_. The number of docked SVs just before the *i* + 1^*th*^ AP is

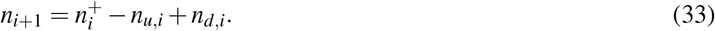

Here *n*_*u*,*i*_ is the number of docked SVs that undock between the *i*^*th*^ and *i* + 1^*th*^ AP, and conditioned on 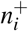 it is given by

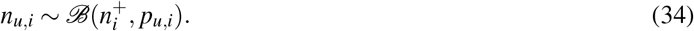

Similarly, the number of empty sites that get occupied in the time interval between successive APs is

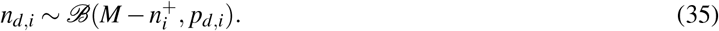

Taking the pgf of *n*_*i*+1_ and using (28)

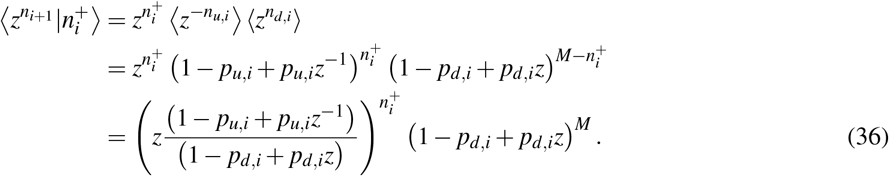

Now unconditioning (36) with respect to 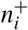 and using (32)

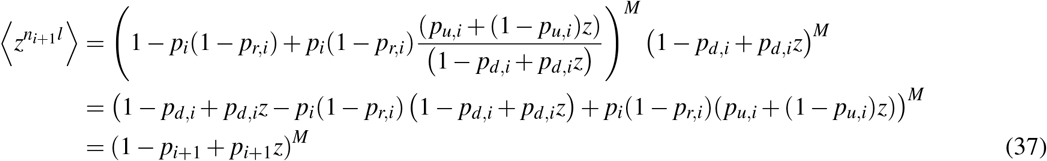

Where

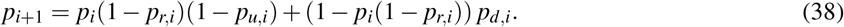

The pgf (37) shows that *n*_*i*+1_ ∼ ℬ (*M, p*_*i*+1_) is a binomially-distributed random variable, hence completing the proof by induction.

## Appendix C

If the time *T*_*i*_ between APs follows an independent and identically distributed (i.i.d.) random variable, then *p*_*u*,*i*_ and *p*_*d*,*i*_ are also i.i.d. random variables drawn as per (25) and (26) in Appendix A, and *p*_*i*_ is the solution to the random discrete-time system (2). In this case, conditioned on *p*_*i*_, *b*_*i*_ is binomially distributed as per (3). However, since now *p*_*i*_ itself is a random variable, *b*_*i*_ will no longer be binomially distributed as has been reported in other works [39]–[41], [45]. For example in [41] we consider AP arrival as per a Poisson process where the inter-spike interval is an exponentially distributed random variable with mean 1*/ f* .Then using the standard tool of moment dynamics for Stochastic Hybrid Systems [70]–[72], the steady-state QC Fano factor *FF* is derived in [41], [67]. Interestingly, in this case, *FF* is maximized at an intermediate frequency *f* (see Fig. 3 in [67]) and for many parameter regimes can be higher than one. Note that the Fano factor of a binomial-distributed random variable is always less than equal to one.

## Appendix D

In this section, we provided an exact analytical solution for the QC autocorrelation function (ACF) that is mathematically defined as the steady-state Pearson correlation coefficient between *b*_*i*_ and *b*_*i*+1_ in limit *i* → ∞.

We first calculate the ACF at lag *𝓁* = 1, conditioned on *b*_*i*_. Thus,

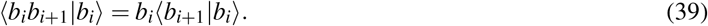

Given *n*_*i*+1_ and the AP-triggered binomial fusion of SVs with probability *p*_*r*,*i*+1_, we can write

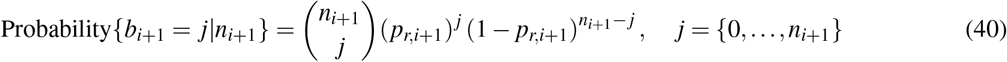

and the mean QC given the number of docked vesicles at stimulus number *i* + 1 follows

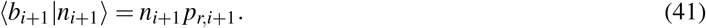

Equation (33) relates the number of docked SVs at *i* + 1^*th*^ AP to those at *i*^*th*^ AP. Thus, given 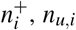 and *n*_*d*,*i*_, *n*_*i*+1_ is known. Rewriting (41)

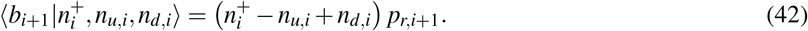

**Fig. S1:**
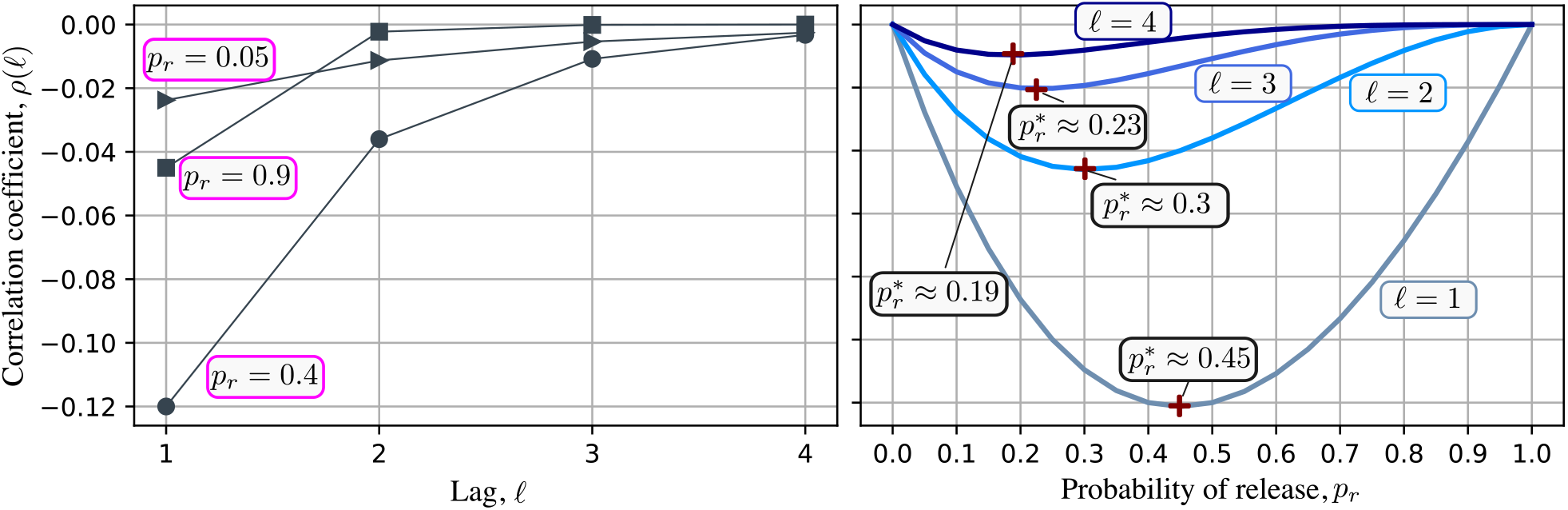
The steady-state correlation coefficient *ρ*(*𝓁*) as a function of lag *𝓁* for different release probabilities with *p*_*d*_ = 0.5 (left). Correlation coefficient *ρ*(*𝓁*) as a function of release probability for different lags *𝓁* and *p*_*d*_ = 0.4.

Hence, by substituting (42) in (39) we obtain

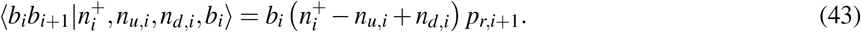

Substituting SVs after AP with those before minus QC yields

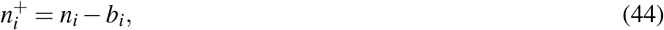

in (43), we obtain

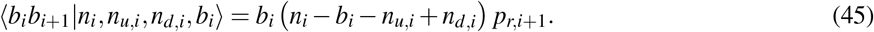

And finally the above expression is simplified to

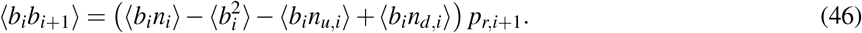

We next calculate ⟨ *b*_*i*_*n*_*i*_⟩, 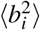, ⟨ *b*_*i*_*n*_*u*,*i*_⟩, ⟨ *b*_*i*_*n*_*d*,*i*_⟩.

*Finding* ⟨ *b*_*i*_*n*_*i*_⟩:

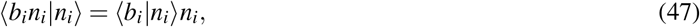

and using (40) for stimulus number *i*

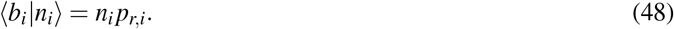

Also using the pmf in (27),

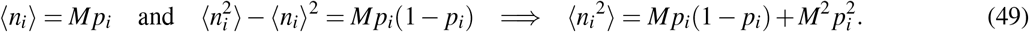

Using (47)-(49)

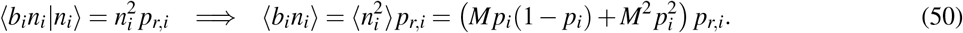

*Finding* 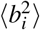:

Using the pmf in (29),

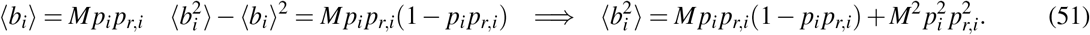

*Finding* ⟨ *b*_*i*_*n*_*u*,*i*_⟩:

Conditioned on 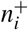 and *b*_*i*_

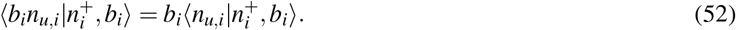

Using the pmf of *n*_*u*,*i*_ (34)

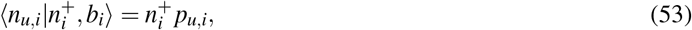

and from (53) and (44)

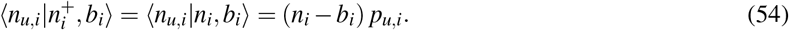

Substituting (54) in (52) we obtain

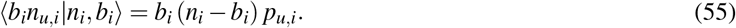

Next, we uncondition the above equation

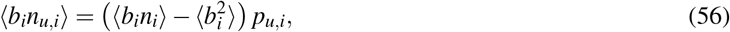

with ⟨ *b*_*i*_*n*_*i*_⟩ is in (50) and 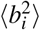 in (51).

*Finding* ⟨ *b*_*i*_*n*_*d*,*i*_⟩:

Given *n*_*i*_ and *b*_*i*_

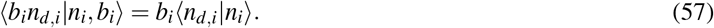

Using (44) and the pmf of *n*_*d*,*i*_ in (35) we can write

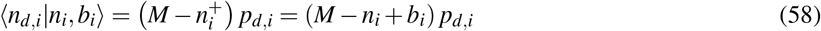

Then we substitute (58) in (57)

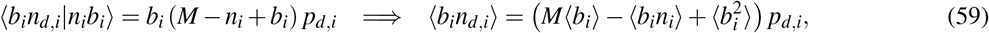

with ⟨ *b*_*i*_⟩ and 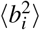 in (51), and ⟨ *b*_*i*_*n*_*i*_⟩ in (50). Finally, we simplify (46) by substituting ⟨ *b*_*i*_*n*_*u*,*i*_⟩ from (56) and ⟨ *b*_*i*_*n*_*d*,*i*_⟩ from (59)

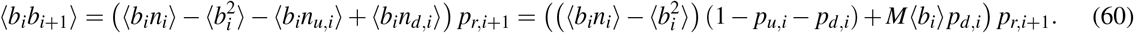

Substituting ⟨ *b*_*i*_⟩ and 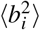 from (51), and ⟨ *b*_*i*_*n*_*i*_⟩ from (50) in (60) we obtain the ACF as follows

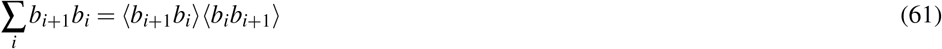

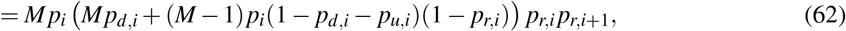

Next we compute the Pearson correlation coefficient for lag *𝓁* = 1 as (18) with ⟨ *b*_*i*_⟩ and 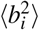 are the mean and second-order moments of QC, respectively. Using constant values of probabilities in (9), (60) simplifies to

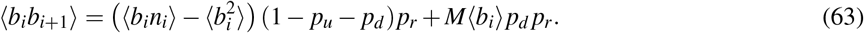

At the steady state

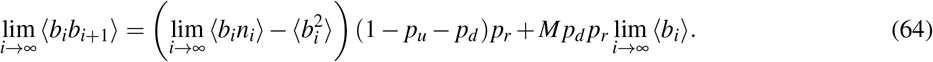

We simplify (50) and (51) for constant values of probabilities and at the steady-state

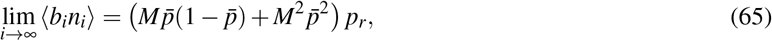

where 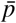 is the steady-state value of *p*_*i*_ (14). Also,

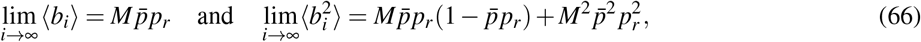

We substitute (12) in (65) and (66) and use them in (64) to obtain ⟨ *b*_*i*_*b*_*i*+1_⟩ at the steady-state as

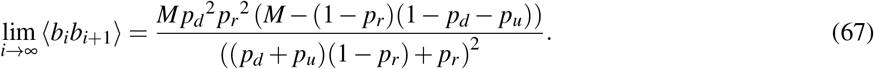

The Pearson correlation coefficient (18) at the steady-state follows

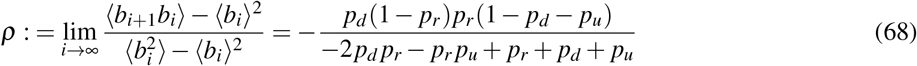

Next, we generalize our approach to calculate the exact analytical solution for the QC autocorrelation function (ACF) and the steady-state Pearson correlation coefficient between ⟨ *b*_*i*_⟩ and ⟨ *b*_*i*+*𝓁*_⟩, where *𝓁* ∈ {1, 2,…}. Assuming constant valued probabilities *p*_*r*_, *p*_*d*_ and *p*_*u*_, the ACF at lag *𝓁* = 1 is calculated in (63). Here we apply the same approach for *𝓁* = 2 following the steps as in (39)-(63). Given *b*_*i*_,

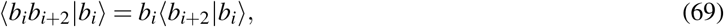

and give *n*_*i+2*_

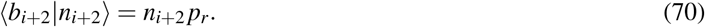

Similar to (33)

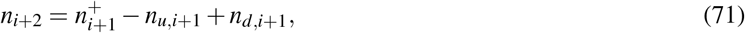

and similar to (44)

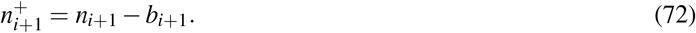

Using (70)-(72) in (69)

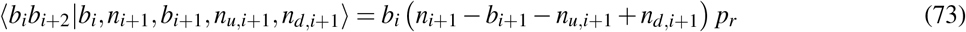

and by unconditioning it we obtain

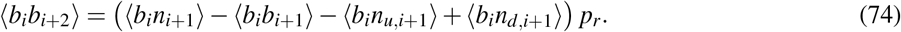

To compute ⟨ *b*_*i*_*b*_*i*+2_⟩ we need ⟨ *b*_*i*_*n*_*i*+1_⟩, ⟨ *b*_*i*_*n*_*u*,*i*+1_⟩ and ⟨ *b*_*i*_*n*_*d*,*i*+1_⟩ which we will find next.

*Finding* ⟨ *b*_*i*_*n*_*i*+1_⟩:

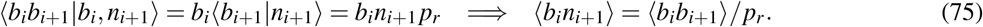

*Finding* ⟨ *b*_*i*_*n*_*u*,*i*+1_⟩:

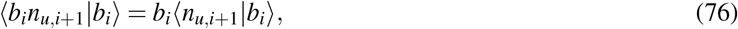

and similar to (54)

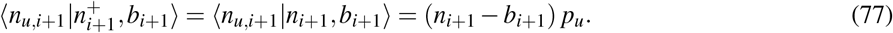

Using (76) and (77)

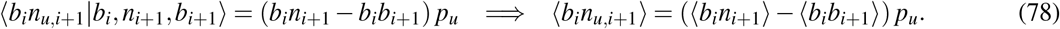

Next, using (75), we simplify the above equation

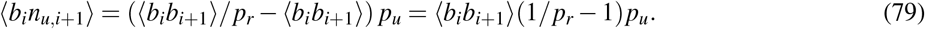

*Finding* ⟨ *b*_*i*_*n*_*d*,*i*+1_⟩:

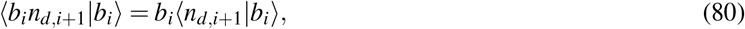

and similar to (58)

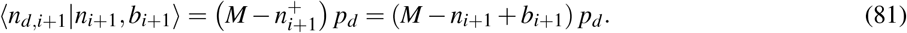

Using (80) and (81) we write

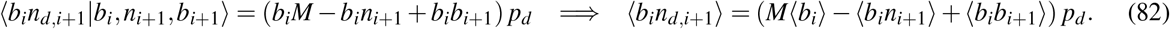

Applying (75) in (82)

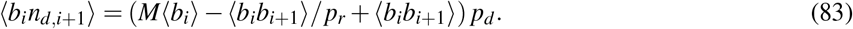

Replacing (75), (79) and (83) in (74) we obtain

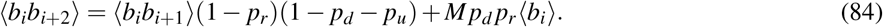

Using (63) in the above equation follows

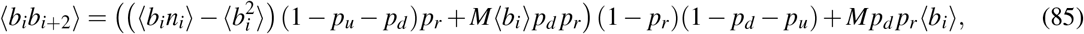

and at the steady-state

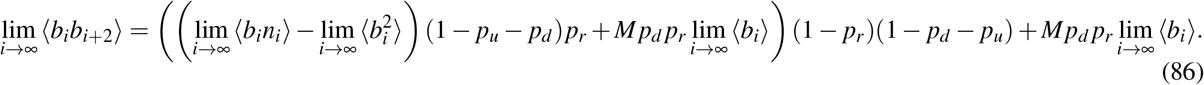

We use (65) and (66) in the above equation when 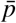 follows (12). Thus,

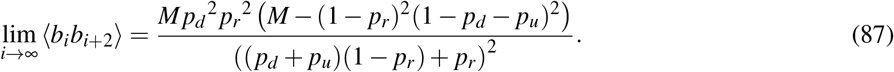

Following the same steps for *𝓁* = 2 in (69)-(87), and by induction we obtain the following expression when

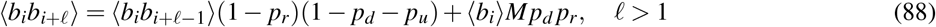

and its steady-state follows

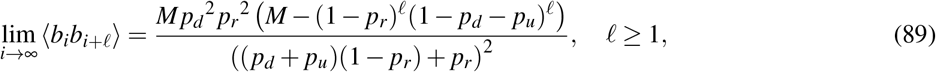

Finally, the steady-state Pearson correlation coefficient follows

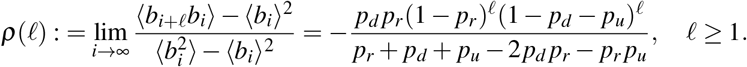

Plots of *ρ*(*𝓁*) as a function of lag *𝓁* and *p*_*r*_ are shown in Fig. S1.

## Appendix E

Of the 19 single-cell recordings obtained from [51], we discarded 3 because of low QCs (QC for first stimulus less than 10), and 1 did not seem to show any synaptic depression. The steady-state QC Fano factor of the 14 single-cell recordings is shown in Fig. S2 as a function of the normalized synaptic depression. To compute steady-state statistics (QC mean and FF) we used stimulus numbers 10 to 800 as some cells began to show a gradual decline in QC after that.

**Fig. S2:**
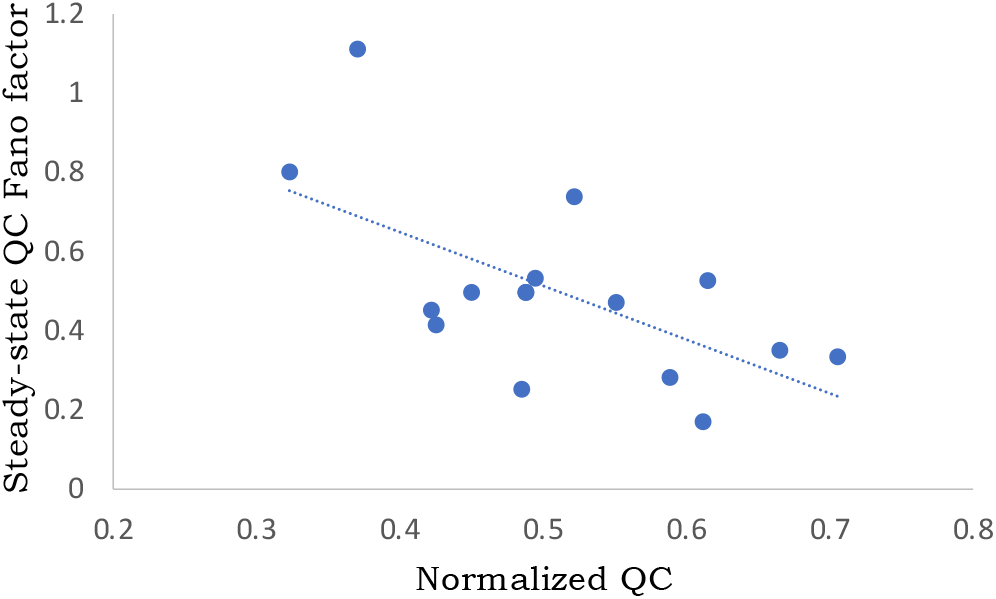
The steady-state QC Fano factor as a function of the normalized QC (steady-state mean QC normalized to the first stimulus) for 14 MNTB-LSO connections from [51]. Each point represents the statistics evaluated from a single-cell QC recording at 50Hz stimulation using stimulus numbers 10 to 800. Consistent with (8), the steady-state QC Fano factor shows a negative correlation with normalized synaptic depression with a Pearson correlation coefficient of −0.6 (*R*^2^ = 0.38), and a 95% confidence interval [−0.85, −0.15].

**Fig. S3:**
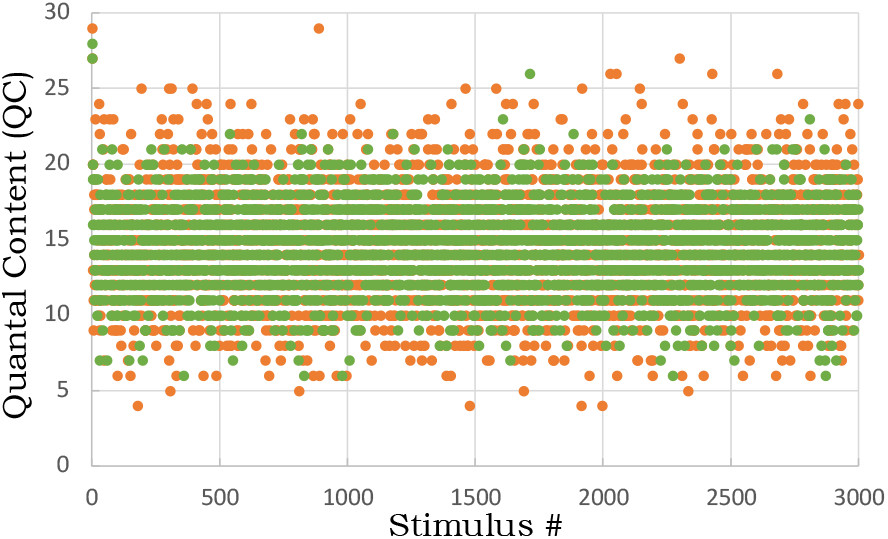
Sample stochastic realizations of the two different models shown in Fig. 6. Orange circles represent the QC per stimulus for the model with low constant probabilities *p*_*r*_ = 0.23 & *p*_*d*_ = 0.2, and the green circles represent QCs for a high release probability *p*_*r*_ = 0.93 and a transient refilling probability as shown in the inset of Fig. 6B that starts high at 0.95 and settles to its steady-state value of *p*_*d*_ = 0.53 after five stimuli.

